# Genome-wide profiling of tRNA modifications reveal the presence of divergent signatures in *Plasmodium falciparum*

**DOI:** 10.1101/2025.02.24.639833

**Authors:** Qian Li, Ylva Veith, Elena Christ, Johan Ankarklev, Ola Larsson, Sherwin Chun-Leung Chan

## Abstract

tRNAs are known to be extensively decorated with various nucleotide modifications, which can modulate the biological properties of tRNAs, such as promoting translation efficiency or fidelity, influencing codon preference and stabilizing tertiary structure of tRNAs. Dynamic alteration of the tRNA modifications, particularly under stress, are conserved gene regulatory mechanisms employed by cells across the kingdoms of life. The malaria-causing *Plasmodium falciparum*, which has a highly AT-biased genome, is reliant on a non-redundant set of tRNA genes to decode distinct and codon-skewed transcriptomes across its developmental stages. This genomic feature highlights the regulatory potential of tRNA modifications and warrants studies on how the tRNA epitranscriptome can impact gene regulatory function in this parasite. However, systematic studies of tRNA modifications in *P. falciparum* remain sporadic. In this study, we present a sequencing-based analysis of tRNA modifications in *P. falciparum* through optimizing the TGIRT-tRNA sequencing approach. The optimized protocol significantly reduces reverse transcription (RT) stop-induced short reads and enhances full-length cDNA synthesis, enabling the identification and stoichiometric estimation of a select nucleotide modifications at single-base resolution. We were able to identify seven types of conserved tRNA modifications and discovered two non-canonical signatures at A59 and U73 positions on specific nuclear-encoded tRNAs, which provides new insights into previously unreported modification sites. Our findings further reveal the dynamic nature of tRNA modifications during stage progression and under stress conditions.

## Introduction

*Plasmodium falciparum* is a protozoan that causes human malaria. The disease continues to impose substantial health burden in Africa and is accounted for an estimated 249 million cases and over 600 000 deaths annually (WHO. 2023). Numerous studies have highlighted the parasite’s adeptness in employing multifaceted strategies to orchestrate tight gene regulation across various developmental stages (Bozdech et al. 2003; Le Roch et al. 2003; Otto et al. 2010). Despite substantial recent progress, these regulatory mechanisms utilized by *P. falciparum* remain incompletely understood. Notably, transcription factor is comparatively underrepresented in the genome and it is consensual that the parasite, unlike other eukaryotes, does not respond to stimulus through an elaborate transcriptional network (Coulson et al. 2004; Balaji et al. 2005), in alternative, post-transcriptional and translational regulatory mechanisms serve important roles in the overall gene regulation, in particularly during cellular stresses (Foth et al. 2008; Foth et al. 2011; Bunnik et al. 2013; Vembar et al. 2016; Hammam et al. 2021b; Small-Saunders et al. 2024a).

*P. falciparum* genome is highly AT-rich and therefore biased in the use of AT-rich codons, yet unlike most of the eukaryotic organisms which possess redundant tRNA genes, it comprises a set of 45 non-redundant nuclear-encoded tRNA genes (Gardner et al. 2002). While the nuclear encoded tRNA genes retain the expected conservation in their sequences and secondary structures shared among tRNAs of eukaryotes (Pütz et al. 2010). The limited tRNAs reservoir is tightly regulated and exploits the imbalance between demand and supply of individual tRNAs to impact differential translation elongation efficiency among transcripts. Since translation elongation has emerged as a prominent gene regulatory mechanism, it is unsurprising that the tRNA epitranscriptome has also been implicated in the regulation of codon-biased translation programme during various processes, such as stage progression and upon artemisinin challenge.

tRNA is the most extensively modified RNA, with an average tRNA molecule containing 8-13 modifications (Zhang et al. 2022). In general, modifications within the anticodon stem loop (ASL) can modulate translational frameshift (Urbonavicius et al. 2001; Gamper et al. 2015; Smith et al. 2022), enhance ribosome binding and specify codon-anticodon recognition (Yarian et al. 2002; Manickam et al. 2016), thus influencing both the efficiency and fidelity of translation. Meanwhile, modifications within the core region of tRNA structure, such as the D loop and the T loop, are essential for maintaining secondary and tertiary structural stability. Emerging evidence have highlighted the dynamic nature of tRNA modification and its significant role in various cellular responses to stimulus (Chan et al. 2010; Chan et al. 2012; Bruch et al. 2020). However, much of these knowledges originated from extensive studies conducted on model organisms such as yeast, mouse and human, but little is known about *P. falciparum*. Nonetheless, a few studies covering this niche did find significant roles of some tRNA modifications, for example stress tolerance in *P. falciparum* is regulated by DNMT2-mediated m^5^C38 methylation in tRNA^Asp^, which seems to be needed for the efficient synthesis of proteins rich in GAC codons (Hammam et al. 2021b). Nonetheless, the roles of other tRNA modifications remain largely unexplored in *P. falciparum*.

Two studies have hitherto conducted a comprehensive survey of tRNA modifications in *P. falciparum* during normal development of the asexual stage and upon artemisinin treatment using liquid chromatography-mass spectrometry (LC-MS/MS)(Ng et al. 2018b; Small-Saunders et al. 2024b). While LC-MS/MS is a powerful technique theoretically capable of detecting all types of tRNA modifications, it lacks the capability to discern the individual tRNAs harboring the identified modifications unless individual tRNAs are isolated from a large amount of cellular RNA. Furthermore, the technique does not give single-base resolution. To overcome these limitations, several alternative techniques have been developed.

One such technique is nanopore sequencing (Smith et al. 2015; Begik et al. 2021; Thomas et al. 2021; Lucas et al. 2023), which offers the advantage of sequencing tRNA directly, bypassing issues related to reverse transcription. However, nanopore sequencing is error-prone and generally performs poorly with short RNA molecules, which is further complicated by the secondary structure of tRNA. Reverse transcription process may still be needed for denaturing the tRNA structure and propelling the tRNA to translocate through the pore (Lucas et al. 2023). Furthermore, the modifications are inferred from the characteristic of electrical current distortion, which is dependent and sensitive to the sequence context and modification type (Begik et al. 2021).

Another method is next-generation sequencing (NGS)-based tRNA sequencing. Since modifications that can directly interfere with the Watson-Crick complementary of a nucleotide base often induces misincorporation or stopping prematurely during reverse transcription (RT) reaction, therefore enabling the identification and quantification of these tRNA modifications at single-base resolution. An added advantage of the method is the simultaneous quantification of tRNA abundance and the estimation of the aminoacylation level. However, the short sequencing reads generated by modifications-induced truncation, such as described in the QuantM-tRNA seq and hydro-tRNAseq protocols (Pinkard et al. 2020), can cause ambiguous multiple alignments during data processing and potential distort the true biological signals. These short reads can also mask the identification of biologically relevant tRNA derived fragments. Therefore, tRNA modification profiling based only on misincorporation, but with minimized truncation, is desired. Several strategies have been developed to overcome this shortcoming, such as using thermostable group II intron reverse transcriptase (TGIRT) to increase processivity and the use of AlkB demethylase to remove certain types of methylation, as described in AlkB-faciliated RNA methylation sequencing (ARM-seq), DM-tRNA-seq, and mim-tRNAseq protocols (Behrens et al. 2021). The highly processive TGIRT also features a unique 3’ template-switching activity which bypass the need of a separate ligation step and reducing issues concerning low adaptor ligation efficiency and ligation bias.

Here, we present a detailed study utilizing TGIRT-tRNAseq, which has been adopted and optimized from mim-tRNAseq, to identify and estimate the stoichiometry of specific modifications in the *P. falciparum* tRNA transcriptome. This method enables the generation of sequencing reads with high coverage and a uniquely mapping ratio. Combining with bisulfite conversion treatment, we were able to detect and quantify seven types of modifications (m^1^A, m^1^G, m^2,2^G, m^3^C, m^5^C, yW and inosine modification) as well as uncharacterized mutation signatures in nuclear-encoded tRNAs. Whereas only m^1^G37 was detected in apicoplast-encoded tRNAs. Our study also demonstrates the applicability of this method for studying the dynamics of tRNA modifications during stage development and under different stress conditions. We anticipate that this work will serve as a valuable reference and resource for future studies on this topic.

## Results

### TGIRT-tRNAseq with low salt condition and low reaction temperature improves tRNA coverage and uniquely mapping ratio

To accurately identify potential modification sites and quantify their stoichiometry in *Plasmodium falciparum*, we adopted and optimized a TGIRT-based tRNA sequencing approach from mim-tRNAseq to profile the tRNA transcriptome across various developmental stages (supplementary figure 1A)(Zheng et al. 2015; Xu et al. 2019). We first focused on improving the generation of full-length cDNA without the use of AlkB demethylase. TGIRT can directly initiate reverse transcription from a 5’ adapter primed on the conserved CCA terminal sequence of mature tRNAs through unique 3’ template switching mechanism, thereby mitigating issues with low ligation efficiency and ligation bias.

Given TGIRT’s versatility across various conditions, as evident from previous reports (Mohr et al. 2013; Behrens et al. 2021), we sought to investigate whether TGIRT could perform optimally under different conditions. To assess this, the small RNA purified from parasites were divided into two groups, one group was treated under standard conditions recommended by the manufacturer (60 °C, 450mM salt), while the other was subjected to both reduced reaction temperature and salt concentration (42 °C, 75mM salt) as suggested by mim-tRNAseq (Behrens et al. 2021). Consistent with their findings, lower reaction temperature and reduced salt concentration (42 °C, 75nM salt) significantly improved tRNA coverage without the aid of AlkB demethylase. Over 80% of aligned tRNA reads spin a complete 5’-3’ coverage, suggesting that this method efficiently mitigate most of the RT stop, enabling the generation of reproducible mutation signatures aligned with frequently modified tRNA positions between biological replicates (Supplementary figure 1B & C). Low salt condition resulted in approximately 80% of reads mapped uniquely to a combined tRNA reference containing *Plasmodium* nuclear and apicoplast tRNAs, as well as human tRNAs (Supplementary figure 1D). Among these, over 90% of the mapped reads originated from *Plasmodium* nuclear tRNAs, with small proportion of reads of human tRNAs origin, while roughly 4% were aligned to apicoplast tRNAs (Supplementary Table 1). Additionally, to improve detection accuracy, we incorporated unique molecular identifiers (UMIs) in the 3’ adapter to minimize PCR duplication bias (Kivioja et al. 2011; Smith et al. 2017).

### Identification and quantification of tRNA methylation sites in *P. falciparum*

Through this approach, we were able to infer and draw an atlas of tRNA modifications in both the nuclear and apicoplast-encoded tRNAs of *P. falciparum*, including m^1^A on A9 and A58, m^1^G on G9 and G37, yW on G37, m^2,2^G on G26, m^3^C on C32 as well as inosine on A34 (I34) (Figure 1A & D). To validate these modifications, RNA samples also treated with *E.coli* AlkB demethylase and its D135S mutant prior to library generation, in line with previous reports, the demethylase treatment significantly lowered the mutation frequency on positions inferred to be modified by m^1^A, m^1^G and m^3^C (Figure 1B)(Cozen et al. 2015; Zheng et al. 2015; Pinkard et al. 2020). However, the demethylase treatment showed very limited effect on m^2^,^2^G.

**Figure 1.**
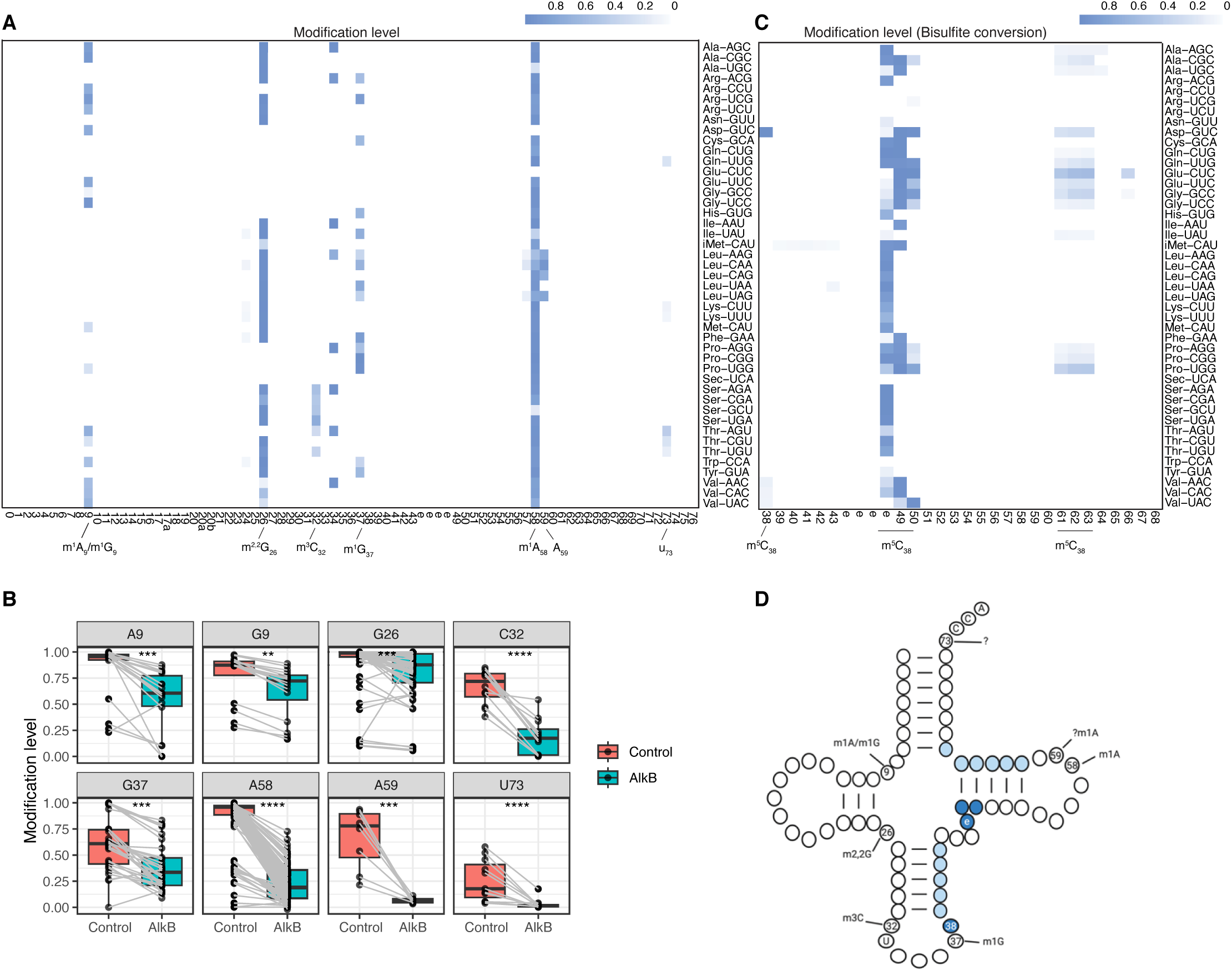
The modification stoichiometry reflected by the mutation frequency observed in low salt TGIRT-tRNAseq. (A) Global heatmap of the average proportion of reads with mutation/truncation signatures detected per position in all tRNA at the ring stage. The numbers represent canonical positions, ‘e’ denotes positions in the variable loop. The inferred identities of the modifications that cause the mutations are shown (A9, G9, G26, C32, A34, G37 and A58); the identities of A59 and U73 cannot be inferred and remain unclear. (N=3 biological replicates) (B) Changes in the modification levels following AlkB demethylase treatment. Modifications on position A59 and U73 were found to be sensitive to AlkB treatment. P-values from Mann-Whitney U-test comparing control and AlkB treatment are denoted by ****(p <0.0001); ***(p<0.001); **(p<0.01). (N=2 biological replicates) (C) Global heatmap of the average proportion of reads with m^5^C methylation detected per position in all tRNA at the ring stage. The numbers indicate canonical positions, ‘e’ denotes positions in the variable loop. (N=3 biological replicates) (D) Schematic representation of the locations of the modifications in relation to the secondary structure of tRNA. Blue represents the presence of m^5^C methylation with high (dark blue) and low (light blue) stoichiometry.

In addition to these conserved modifications, we also observed unique mutation signatures at previously unreported sites, such as at A59 of all tRNA^Leu^ except tRNA^Leu-UAA^ as well as at the discriminator base U73 of all tRNA^Thr^, tRNA^Lys^ and tRNA^Gln-UUG^. Although the identities of these signatures remain unclear, as no relevant annotations have been reported or deposited in MODOMICS (Cappannini et al. 2024), both A59 and U73 signatures were sensitive to demethylase treatment and thus likely to be methylations. This suggests that A59 may also be methylated on N1 position similarly as on A58 (N^1^-methyladenosine, m^1^A) (Figure 1B). Since modifications at U73 can give rise to mutation signature, we hypothesize this modification may directly disrupt Watson-Crick pairing and making it likely to be m^3^U modification. However, further investigation is required to confirm the modification type at these two positions.

The proportion of misincorporation and/or RT stops ratio is widely utilized as a method to quantify modification levels. Notably, our method significantly improves the generation of full-length cDNA, enabling near-complete modification readthrough for most of the detected modification types, except for m1A58, where the juxtaposed modifications at A58 and A59 in tRNA^Leu^ induce deletions. Our optimization thus improves the accuracy in estimating modification levels (Supplementary figure 2). To better review the tRNA epitranscriptome, we categorized the modifications into three groups base on their estimated stoichiometry (low: 5%-40%, hypomodified; moderate: 40%-70%; high: 70%-100%, hypermodified) (Figure 1A and Table 1).

m^1^A58 is recognized for its pivotal role in maintaining the proper structure of tRNA^iMet^. The presence of m^1^A58 can drastically increase the thermostability of the tRNA that harbors it (Droogmans et al. 2003), while its absence promotes tRNA degradation via nuclear surveillance and RTD pathways (Kadaba et al. 2004; Tasak and Phizicky 2022). As expected, 40 out of 45 tRNAs exhibit high levels of methylation. Consistent with previous reports on other organisms, tRNA^Asp(GUC)^ and tRNA^Glu(CUC)^ are hypomodifed (below the 5% threshold) (Kuchino et al. 1981; Clark et al. 2016; Behrens et al. 2021). In addition, tRNA^Ala(UGC)^, tRNA^Ile(UAU)^ and tRNA^Ser(GCU)^ also show low modification levels in *P. falciparum*, in which the latter two contrast findings in other organisms.

m^2^G26/ m^2,2^G26 is another modification that is crucial for maintaining proper tRNA structure and stability. The singly or doubly deposited methyl-groups on N2 occurs in the hinge of D-loop and the anticodon loop and has been proposed to prevent the mispairing with C11 and therefore ensuring the integrity of the tRNA tertiary structure (Pallan et al. 2008). We found this modification is universal and hypermodified among all the nuclear tRNAs in *P. falciparum*, so much if a tRNA retains the G26 identity. Although both m^2^G and m^2,2^G can be found on G26, the mutation signatures likely arise from the di-methylated base because the mono-methylated exocyclic nitrogen is not expected to disrupt the Watson-Crick pairing. Interestingly, the stoichiometries of m^2^G26/ m^2,2^G26 on all tRNA^val^ are significantly lower than other tRNAs. Notably, only tRNA^val^ have U11 instead of C11 among all the tRNAs that have G26 identity. This divergence may have reduced the propensity of mispairing between unmodified G26 with position 11 and alleviated the stringent need of modification on this position.

Adenosine to Inosine editing on the wobble position of ANN anticodon (I34) within all the four-codon family boxes are important mechanism for expanding the decoding capacity of these tRNAs and for achieving efficient translation of proteins. (Torres et al. 2021). I34 is detected in all eight tRNAs that are known to undergo A-to-I editing in eukaryotes, with most of these tRNAs found to be completely edited, corroborating the ubiquitous nature of I34 on these target tRNAs. However, inosine modification at position 37 (I37), which is commonly found in eukaryotic tRNA^Ala^ (Gerber et al. 1998), was not detected in in *Plasmodium falciparum*.

Unlike m^1^A58, m^2^G26 and I34, which are often hypermodified in tRNAs that harbor the modifications, the modification levels of m^1^A9/m^1^G9, m^3^C32 and m^1^G37 vary substantially among individual tRNAs.

m^1^A9/m^1^G9 are modifications on position 9. Their presences may be important for the correct folding of tRNA into the cloverleaf structure (Helm et al. 1998; Sakurai et al. 2005). In eukaryotes, m^1^G9 is prevalent in cytosolic tRNAs, while m^1^A9 is mostly restricted to mitochondrial tRNAs (Suzuki 2014; Vilardo et al. 2020), with the exception of cytosolic tRNA^Asp(GUC)^ (Clark et al. 2016). Interestingly, we could detect m^1^A9 in 9 cytosolic tRNAs, including tRNA^Asp(GUC)^, which are often hypermodified. Meanwhile, m^1^G9 is found on 8 tRNAs, albeit mostly moderately modified. Although, the methylase that catalyzes methylation on position 9 of mitochondrial tRNAs is reportedly bifunctionally active towards both A and G (Vilardo et al. 2012), the unique status of m^1^A9 modification in *P. falciparum* cytosolic tRNAs may represent a targetable process.

Methylation of guanosine N1 position on the immediate 3’-side of the anticodon (m^1^G37), is a highly conserved modification and can contribute to the faithful maintenance of the reading frame by suppressing ribosomal +1 frameshift error. m^1^G37 is also reportedly coupled with efficient aminoacylation of the underlying tRNAs and is essential for cell viability (Gamper et al. 2015; Clifton et al. 2021). m^1^G37 is observed in 15 cytosolic tRNAs decoding 7 different amino acids, which notably represent all that have a G37 identity, suggesting a non-selective activity by the responsible RNA methyltransferase. However, as expected of a regulatory function, the modification levels among these tRNAs display a wide range of variation. Furthermore, only some members of tRNA^Arg^, tRNA^Ile^ and tRNA^leu^ are m^1^G37 modified, the differential status of this modification among isoacceptors may contribute to codon-biased translation regulation.

Wybutosine (yW) is a complex modification found exclusively at position G37 of both eukaryotes and archaeal tRNA^Phe^ (Perche-Letuvée et al. 2014). Despite both yW and m^1^G37 give rise to mutation signatures, previous characterization provides a basis to discern mutation signatures derived from yW or m^1^G37 modification. It was observed that yW mostly caused misincorporation of thymidine (T), whereas m^1^G37 modification causes cytosine misincorporation during RT reaction (Behrens et al. 2021). The mutation pattern we observed on tRNA^Phe(GAA)^ also differs from the typical m^1^G37 mutation signature and is consistent with previous study. Therefore, it is expectedly that *Plasmodium* tRNA^Phe^ also harbors yW modification instead of m^1^G at position 37. However, the G37 mutation signature on tRNA^Arg(UCG)^ also distinctly misincorporated adenosine at high frequency, suggesting that the pattern of mutation signature can be dependent on the sequence context (Supplementary figure 3).

m^3^C modification on the 5’-side of the anticodon (m^3^C32), is conserved in eukaryotes and is commonly found in tRNA^Thr^, tRNA^Ser^ and tRNA^Arg^. The function of m^3^C32, however, remains debated. While it is expected to regulate translation given its proximity to the anticodon, deletion of its methyltransferases is viable and resulted in very mild phenotype. Although studies have also found codon-specific translation impairment upon in m^3^C32 methyltransferase-deficient cells, in particularly for mitochondrial translation (Arimbasseri et al. 2016; Kleiber et al. 2022). m^3^C can be found in the variable loop of tRNA^Ser^ as well as at position 20 of tRNA^Met^ in eukaryotes (Clark et al. 2016; Behrens et al. 2021; Cui et al. 2021). In *P. falciparum*, we observed mutation signatures at C32 only in tRNA^Ser^ and two of the three RNA^Thr^, ranging from low to moderate stoichiometry. Whereas there is no indication of m^3^C modification at position 20 and 47 in any of the tRNAs.

In addition to conserved modifications that can be inferred from the mutation signatures. We also observed unexpected signatures at A59 and U73. A59 signatures are exclusively found in four of the five tRNA^Leu^, which manifest mainly as misincorporation but also result in deletions spanning position 57 to 59. With the exception of tRNA^Leu(UAA)^, which devoid of A59 signature despite retaining the identity, the other four tRNA^Leu^ are hypermodified at this site (Supplementary figure 4). Meanwhile, U73 mutation signatures are detected in all tRNA^Thr^ and tRNA^Lys^ as well as tRNA^Gln(UUG)^. Position 73 often serves as a crucial discriminator base for accurate and efficient aminoacylation of the tRNAs by the cognate tRNA synthetases, therefore, modification on U73 may influence this process and the outcome of the translatome. Furthermore, the stoichiometry is relatively low for most U73 signatures, a feature that shares more resemblance to modifcations located within the anticodon loop, than the structurally related modifications (Supplementary figure 5).

Since A59 and U73 signatures are sensitive to demethylase treatment, we believe the signatures may correspond respectively to m^1^A and m^3^U modifications.

While the same type of modification is often found on all isoacceptors decoding a specific amino acid, some isoacceptors harbor different modification patterns among themselves. For example, tRNA^Thr^, tRNA^Arg^ and tRNA^Ala^ differ among themselves in multiple positions, these disparities may subject isoacceptors to differential posttranscriptional regulation.

In addition to the standard sequencing, we also performed bisulfite (BS) conversion prior to tRNA library construction, which relies on the addition of sodium bisulfite to deaminate all unmodified cytosines into uracils while leaving m^5^C modified cytosines unaffected, thus the level of m^5^C modifications (Schaefer et al. 2009). Positions that are heavily deposited with m^5^C include the conserved C38 of tRNA^Asp-GUC^ and the nucleotides within the variable loop of most tRNAs. Notably, clusters of m^5^C are also observed on both sides of the T-stem of many tRNAs. Although observed in low stoichiometry, the side that juxtaposes the acceptor stem is not known to harbor m^5^C in other organisms to the best of our knowledge (Figure 1C & D).

As a test of the applicability of our protocol, we generated a m^3^C32 deficient strain using the previously described CRISPR-Cas9 editing to delete the *P. falciparum* tRNA N(3)-methylcytidine methyltransferase ortholog, PF3D7_1248100 (Ghorbal et al. 2014). The KO strain did not show any growth or visible morphological phenotypes. We assessed the functionality of this gene by comparing the mutation signature profiles between the KO strain and the wide-type NF54 strain. As anticipated, only the C32 positions were completely devoid of any mutation signatures, while no other comparable changes in the profile were observed (Supplementary figure 6). Therefore, PF3D7_1248100 is the sole methyltransferase responsible for m^3^C32 methylation and it is non-essential during asexual development.

Taken together, our optimized sequencing protocol and analysis pipeline can accurately reflect the modification stoichiometry in single-base resolution.

### Modifications identified in the apicoplast-encoded tRNAs

Among all the mapped tRNA reads, approximately 4% were exclusively aligned to apicoplast tRNAs (Supplementary Table 1). Unlike the mitochondrial genome of *P. falciparum,* which does not encode any tRNA gene, the 33kb apicoplast genome encodes a total of 33 tRNA genes that transcribe 26 unique tRNAs (with seven of them duplicated due to a single inverted duplication event), this minimum anticodon repertoire necessitates superwobbling during decoding (Gardner et al. 2002). Similar to their nuclear counterparts, tRNAs containing G at position 37 were all modified with medium to high level of m^1^G, and surprisingly, this represents the only modification in apicoplast tRNAs detectable by tRNAseq (Figure 2A).

**Figure 2.**
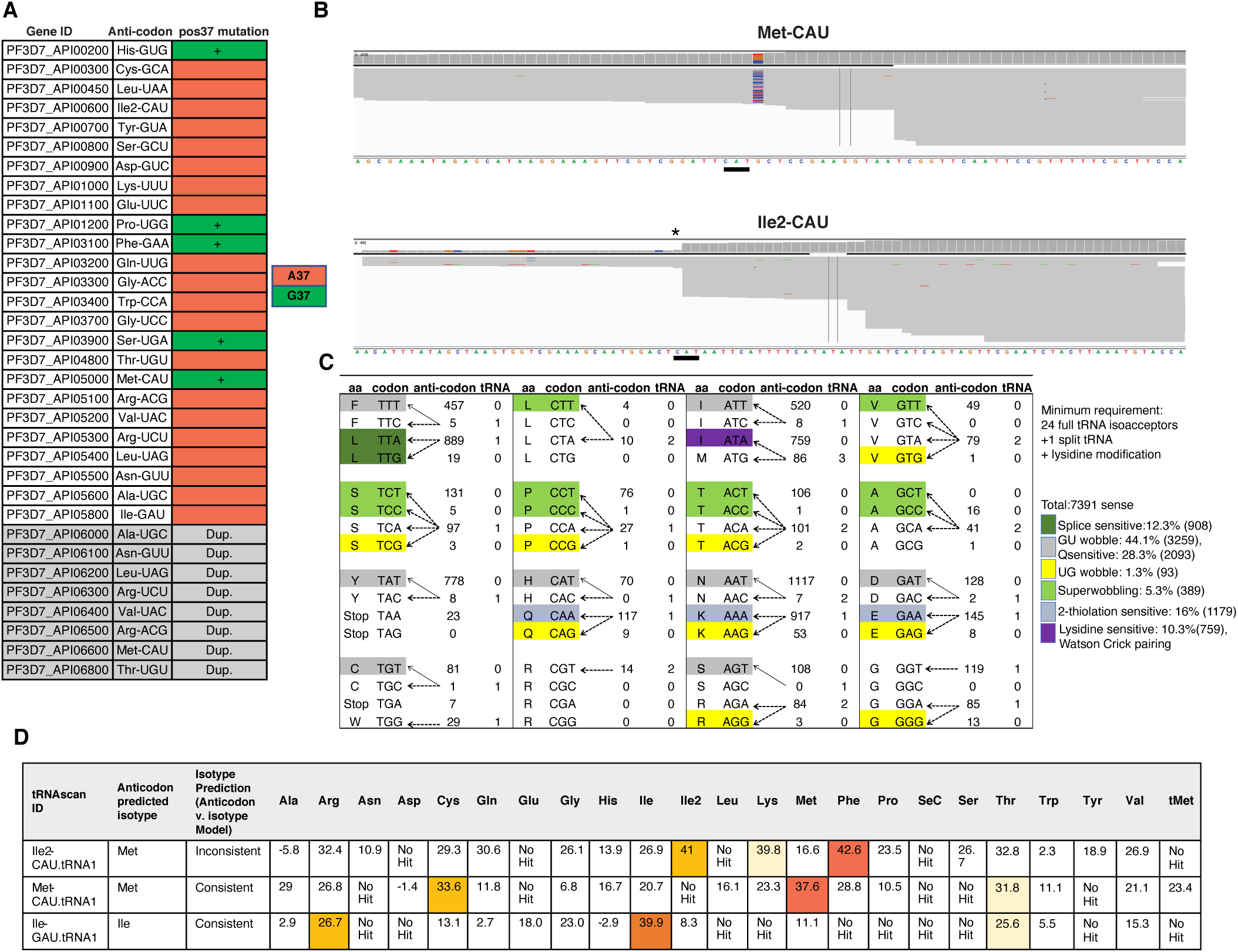
The characteristics of the apicoplast tRNA epitranscriptome. (A) Table of methylation status of apicoplast tRNA detected by low salt TGIRT-tRNAseq. Only m^1^G37 is detected but not the highly conserved m^1^A58. All G37 sites on apicoplast tRNAs are methylated. Nucleotide identity on position 37 is denoted by A37 (orange) and G37 (green). (+) denotes detected mutation signature. (Dup.) indicates duplicated genes. (B) Integrative Genomics Viewer (IGV) shows the truncation on the wobble position (*) of a potential Ile2-tRNA (PF3D7_API00600, lower panel), with no truncation observed in the tRNA^Met^ (PF3D7_API05000, upper panel). (C) An illustration of all possible codon-anticodon pairings predicted by the wobble hypothesis. The cognate tRNA for Ile-ATA codon is not found in the apicoplast genome. The number of tRNA genes per anti-codon (tRNA) and the number of codons used (codon usage) in the apicoplast genome are indicated. (D) Prediction of PF3D7_API00600 as tRNA^Ile2^ by tRNAscan-SE.

Furthermore, we noticed a strong truncation and mutation signature aligned to the wobble position in one of the tRNA^Met^ (PF3D7_API00600, Figure 2B). Using tRNAscan-SE (Chan and Lowe 2019), the tRNA was predicted as an Ile2-tRNA (Figure 2D). Since the cognate tRNA for decoding the frequently used Ile-AUA codon is absent in the apicoplast genome (Figure 2C), we deduce that the signatures should correspond to lysidine modification (K^2^C), which is the addition of a lysine moiety on the cytosine of the wobble position of a tRNA^Met^ to convert the codon specificity of the underlying tRNA from decoding AUG(Met) to decoding AUA(Ile) (Muramatsu et al. 1988; Marck and Grosjean 2002). Notably, while lysidine modification is prevalent in bacteria, it is only sporadically reported in eukaryotic organelle. Therefore, lysidine modification may represent a targetable process.

### Modification level during stage transition

tRNA epitranscriptome can directly and indirectly affect global mRNA translation and impart biological significance. We sought to assess whether these modifications can be dynamically modulated when the parasite develops within the host red blood cell. We isolated tRNA from tightly synchronized cultures at three distinct developmental stages across the asexual replicative cycle: ring (8-12 hours post invasion, h.p.i.), trophozoite (24-28 h.p.i.) and schizont (40-44 h.p.i.) Notably, we only observed a modest reduction in global modifications in trophozoite stage (Figure 3A). Trophozoite stage is known for significant spike in transcriptional and translational activities, we hypothesize the global reduction in tRNA modifications may be correlated with the increased transcription of nascent tRNA which create a backlog of hypomodified tRNAs pending for further processing. To clarify this trend of transient reduction, we specifically examined tRNA^Tyr-GUA^.

**Figure 3.**
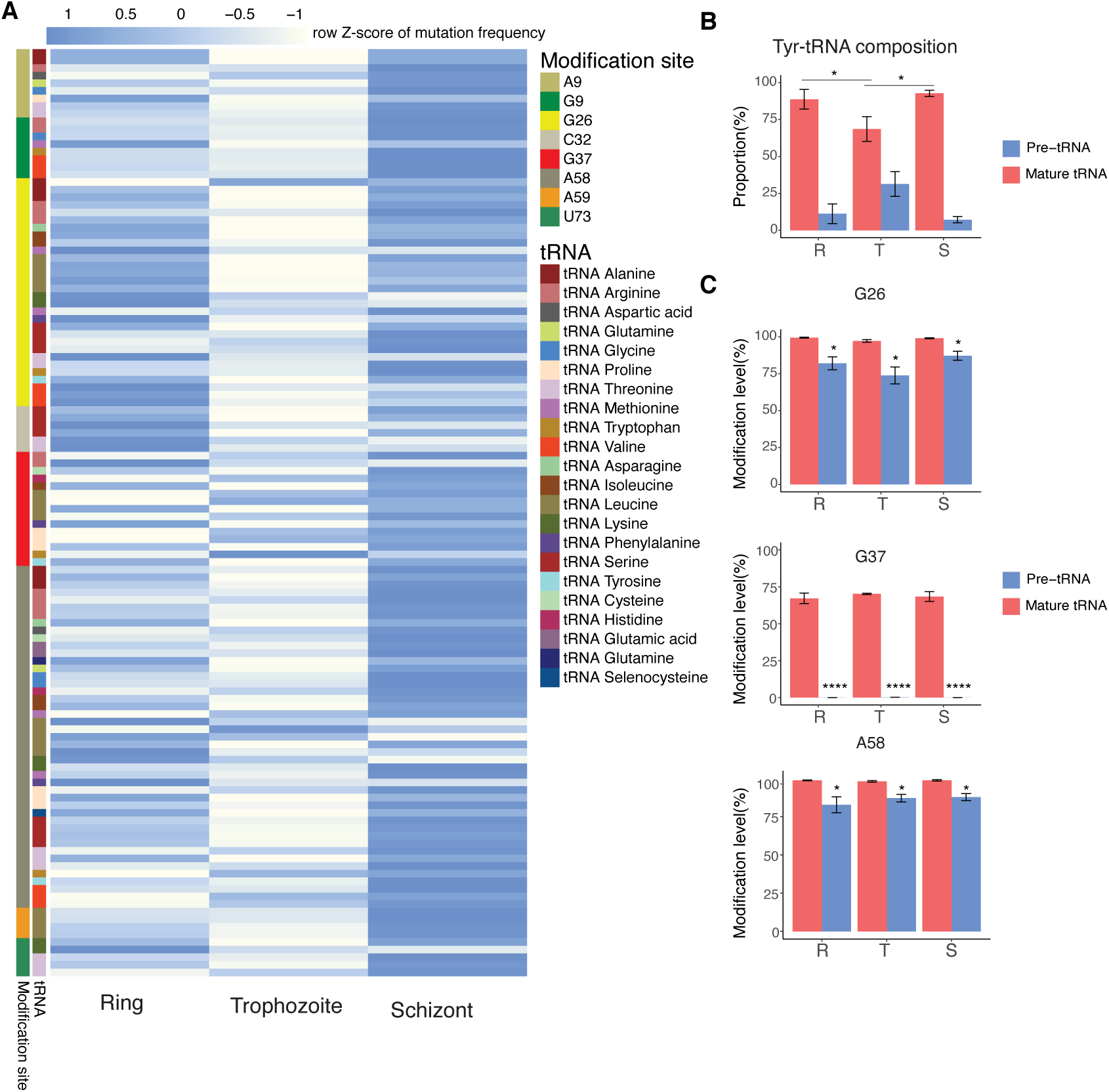
The dynamics of tRNA modifications throughout the IDC. (A) Heatmap illustrating changes in the average modification levels at each site during IDC. (N=3 biological replicates in each stage) (B) The relative proportion of pre- and mature tRNA^Tyr^ during the IDC. P-values from a two-tailed unpaired t-test comparing ring stage and trophozoite stage mature tRNA^Tyr^ or comparing trophozoite stage and schizont stage mature tRNA^Tyr^ are denoted as *(p<0.05) Error bar represent the standard deviation of N=3 biological replicates (C) Mutation frequencies of pre- and mature tRNA^Tyr^ at G26, G37 and A58 modified sites during the IDC. P-values from a two tailed unpaired t-test comparing modification level between pre-mature and mature tRNA^Tyr^ at each modified site are denoted as ****(p<0.0001); *(p<0.05). Error bars represent the standard deviation of N=3 biological replicates

Since it is the only tRNA that splices an 11-nt intron from its pre-tRNA, the ratio of unspliced to spliced tRNA reads can be used as a proxy for nascent tRNAs transcription. Expectedly, trophozoites express a higher level of unspliced tRNA when compared to ring and schizont stages (Figure 3B). Furthermore, the unspliced tRNAs harbor lower levels of modifications at all sites compared to the mature spliced tRNAs, especially with m^1^G37 which is completely absent (Figure 3C). Therefore, the global reduction observed in tRNA modifications is likely due to the increased transcription of pre-tRNAs during the transcriptionally active trophozoites stage rather than as an *ad hoc* regulatory mechanism. Our result, therefore, does not suggest dynamic regulation of the detectable modifications during the asexual cycle, Gametocytes represent the sexually committed or differentiated stage of the parasite, which is the naturally transmissible stage from the human host. Gametocytes exhibit drastic difference in the expression patterns of genes and proteins when compared to the asexual stages. This substantial biological difference prompts us to investigate the dynamics of the tRNA epitranscriptome during gametocytogenesis.

During the process of sexual conversion, the parasites undergo a series of developmental transition over a period of about 8-12 days, moving through 5 distinct morphological phases (designated as stages I-V) to reach the fully differentiated stage V gametocytes. To investigate the change in tRNA modifications during stage I-IV of gametocyte development, *P. falciparum* NF54-PfPeg4-tdTomato parasites were induced to undergo gametocytogenesis and monitored for the next 7 days to recapitulate the process. NF54-PfPeg4-tdTomato has a genomic integrated tdTomato gene under the control of gametocytogenesis-specific promoter, therefore, the process can be monitored in real-time (McLean et al. 2019). Parasites collected at Day-1 prior to induction, Day1, Day3, Day5 and Day7 post-induction were collected. While most tRNA modification sites maintained a steady level during gametocyte commitment, we observed a consistent increase in the modification levels of m^1^A9, m^2,2^G26, m^1^A58 and U73 in some specific tRNAs from the trophozoite stage to Day3 gametocytes. Notably, some of the U73 mutation signatures showed nearly threefold increase. These increases were subsequently stabilized or reverted to slightly decreased from Day3 onwards (Figure 4A-D). Interestingly, most of these modifications are outside the anticodon loop and are important for maintaining structural integrity and stability.

**Figure 4.**
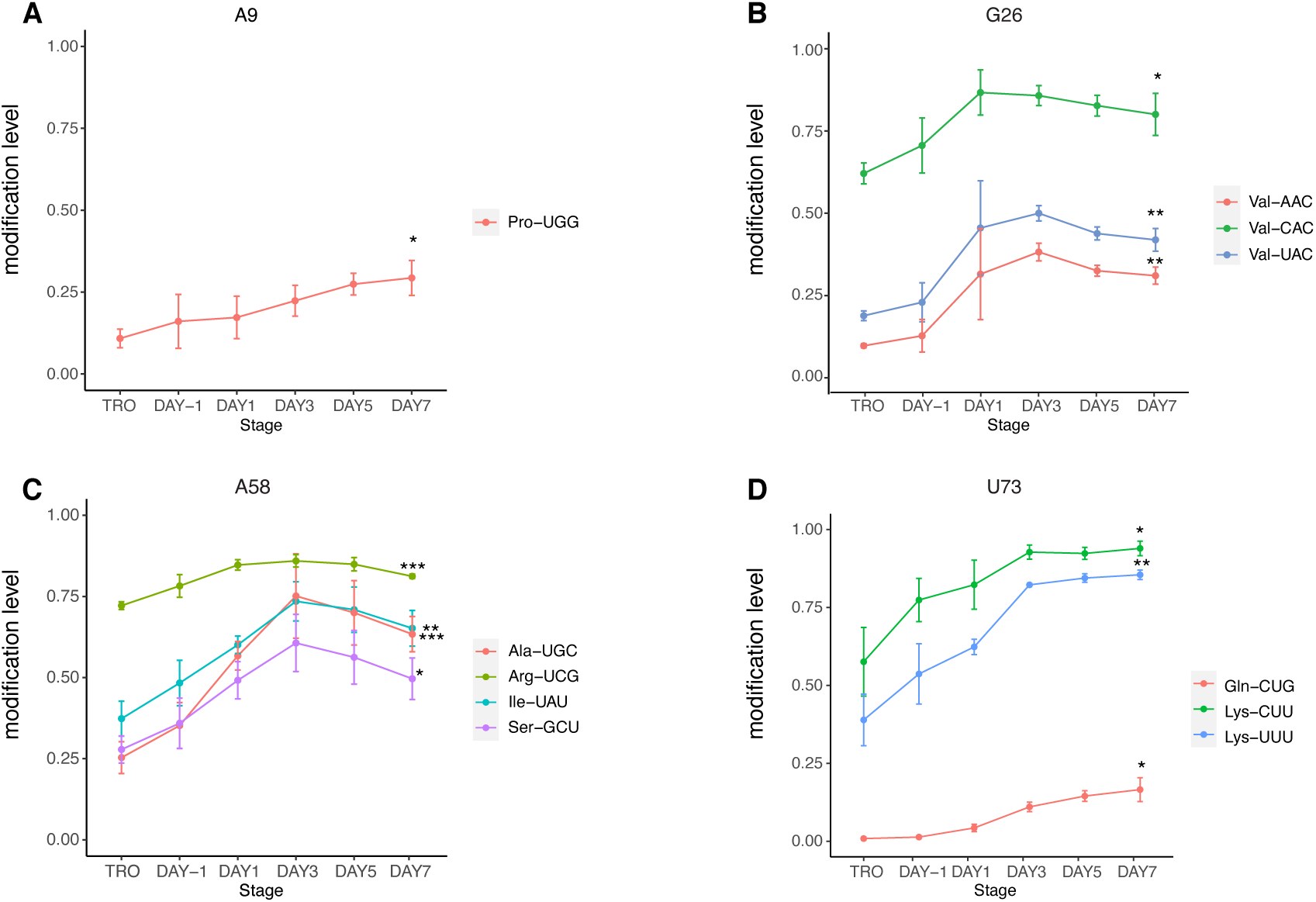
The changes in tRNA modifications during the course of sexual differentiation. (A-D) The changes in modification levels of specific tRNAs during the transition from the asexual stage (trophozoite) to the sexual stage at modified sites A9, G26, A58 and U73. Day-1 to Day7 correspond to the numbers of days post-gametocyte induction. P-values from a two tailed unpaired t-test comparing modification level between trophozoite stage and DAY7 gametocyte are denoted as ***(p<0.001); **(p<0.01); *(p<0.05). Error bars represent the standard deviation of N=3 biological replicates.

### Artemisinin treatment results in an increase in tRNA modifications

Mounting reports have implicated a role of the tRNA epitranscriptome in stress response in bacteria and yeast as well as in the progression of human diseases (Dedon and Begley 2014; Gu et al. 2014; Endres et al. 2015; Zhang et al. 2022). How this may be relevant in *Plasmodium spp.,* however, is rarely considered. A recent study showed that U34 thiolation can reprogram the translatome through codon-biased translation and contributes to artemisinin resistance in *P. falciparum*, in particular, U34 thiolation elevated the synthesis of Kelch13 proteins, which is the consensual resistance marker (Small-Saunders et al. 2024b). This represents the first study that mechanistically link the tRNA epitranscriptome to stress response.

To further investigate the role of tRNA modifications in *Plasmodium* stress response, we subjected parasites to different stress conditions including amino acid starvation, glucose starvation and artemisinin treatment.

Despite this effort, we did not observe any notable alterations in tRNA modification levels upon short-term amino acid starvation or glucose starvation (data not shown). However, when tightly synchronized early ring stage parasites (0-6 hpi) were treated with 700nM DHA (dihydroartemisinin) or DMSO for 6 hours, we observed significant differences (Figure 5A). We found that most of the detectable tRNA modifications were slightly elevated in the DHA-treated group when compared to the DMSO-treated group. Furthermore, the elevation, despite small, shows dose dependency on DHA concentrations. Among the changes, the modifications on A59 and U73 showed the most noticeable increases, with at least a 2-fold increase in the stoichiometry upon DHA treatment (Figure 5B).

**Figure 5.**
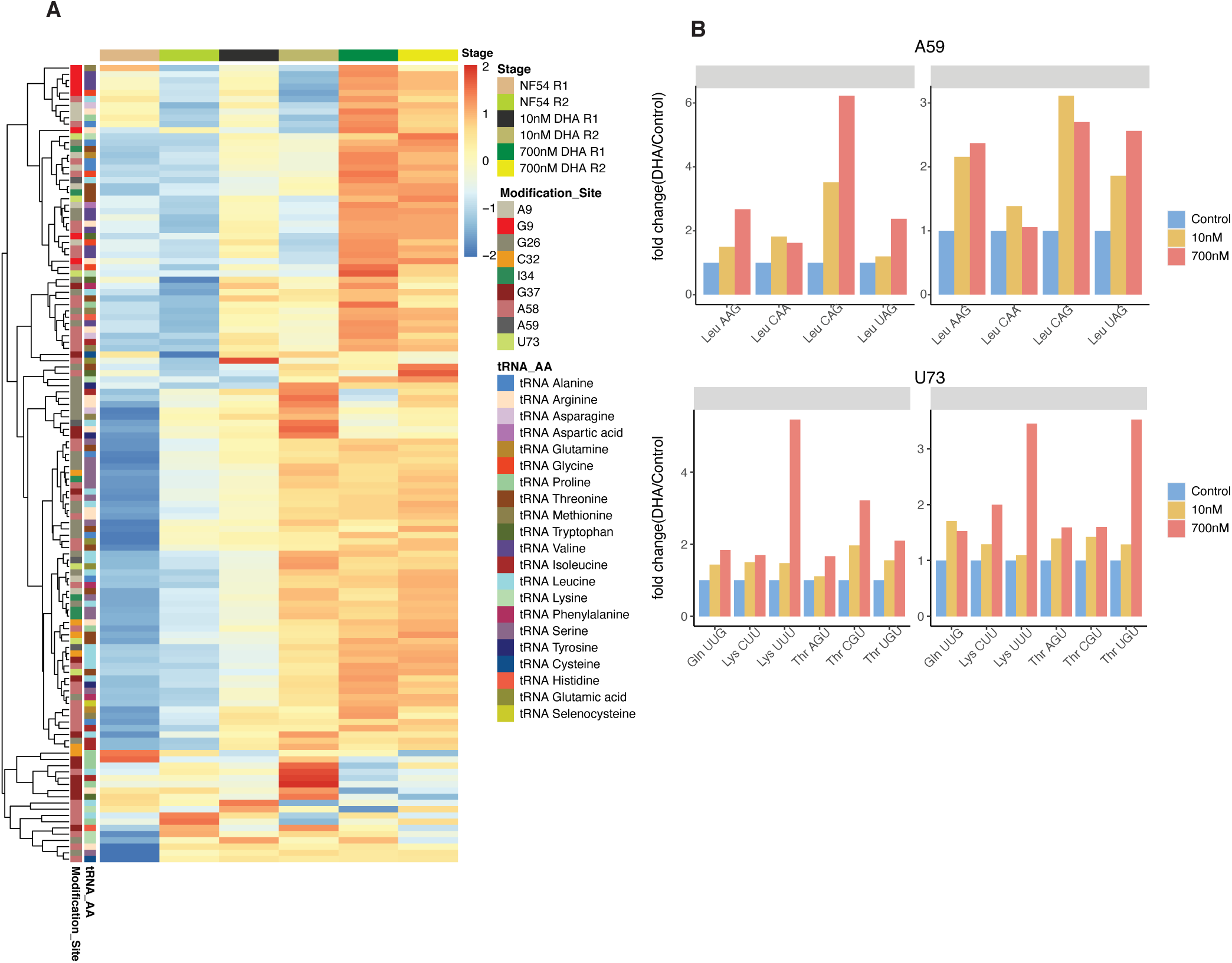
The changes in tRNA modifications upon DHA treatment. (A) Heatmap illustrating the changes of modification levels under DHA treatment (N=2 biological replicates in each condition). (B) The fold change of modification levels at position A59 and U73 under DHA treatment.

## Discussion

Here, we present the characterization of tRNA modifications in the malaria parasite *P. falciparum* at single-base resolution using tRNAseq. Together with previously characterization of tRNA modifications using mass spectrometry (MS) (Ng et al. 2018b), we complement the landscape of the tRNA epitranscriptome in *P. falciparum* which can serve as a reference for later studies on this topic.

By reducing the salt concentration and reaction temperature, we could significantly increase the yield of full-length cDNA and improved the read coverage in comparison to the standard TGIRT protocol, allowing us to identify and quantify tRNA modifications in *P. falciparum*.

Through the generation of PF3D7_1248100 knockout strain, we have validated that PF3D7_1248100 is solely responsible for catalyzing N(3)-methylcytidine methylation on tRNA as the knockout strain resulted in the complete depletion of m^3^C32 modifications on both tRNA^Ser^ and tRNA^Thr^, underscoring the robustness and sensitivity of our method.

In addition to the conserved modifications that are frequently reported with tRNAseq, we have identified two mutation signatures in A59 and U73, two previously unreported positions. While further work is required to confirm the identities of these modifications, they are sensitive to AlkB demethylase treatment and would be suggestive of methylations at both sites. Since most of the mutation signatures arise due to the presence of modifications that can directly interfere with the Watson-Crick pairing, therefore, we deduce that A59 signatures can either be m1A or m6,6A modification. However, m6,6A modification is hitherto found to be restricted to rRNAs, pointing convincingly to an m1A identity at A59. In archaea, position A57 of tRNAs are also methylated by the same methyltransferase that catalyzes m1A58 modification in a sequence context dependent manner, though the functional significance of A57 modification remains unknown (Hamdane et al. 2014). It may be possible that A59 is similarly methylated by a bi-specific A58 methyltransferase.

Meanwhile, we deduce that the U73 signatures most likely reflect m^3^U modifications. While m^3^U modification is also known to be restricted to rRNAs (Patrasso et al. 2023), a previous MS study had detected m^3^U in purified *P. falciparum* tRNAs (Ng et al. 2018b), the discovery is hitherto unique in this species and corroborates our current deduction (Cappannini et al. 2024). U73 modifications are located just outside the the acceptor stem of tRNA^Thr^ and tRNA^Lys^, adjacent to the post-transcriptionally attached -CCA tail. This position is commonly referred as the discriminator base of a tRNA and is important for the addition of the -CCA tail as well as ensuring the fidelity and efficacy of the aminoacylation process (Wende et al. 2015). For instance, mutations at C38 and G73 alone in a yeast tRNA^Arg^ is capable of converting its specificity into an Asp acceptor (Fender et al. 2004); a U73 substitution in *E. coli* tRNA^Cys^ can abolish recognition and charging by cysteine tRNA synthetase (Hamann and Hou 1995), while C73U mutation of tRNA^His^ significantly reduces aminoacyl transfer rate (Bovee et al. 1999). Interestingly, we also observed an unexpectedly low equilibrium aminoacylation level in *P. falciparum* tRNA^Thr^ (Li et al. 2024), prompting our speculation that modification at U73 may affect the aminoacylation process. U73 modifications were significantly increased during early sexual commitment and upon exposure to DHA stress. Since trigger of the *in vitro* gametocytogenesis process involves nutrient stresses, the convergent trends suggest potential regulatory role of this modification in response to certain stress conditions.

The previous MS study did not detect any yW modification even though the genome of *P. falciparum* encodes all of the processing enzymes necessary for yW biosynthesis (Iyer et al. 2010; Sawhney et al. 2015; Ng et al. 2018a). However, in our study, the mutation signatures found on the only known position, G37 of tRNA^Phe^, closely resemble the pattern reported in the analysis of the mim-tRNAseq data, in which adenosine misincorporation was overrepresented (Behrens et al. 2021). This infers the presence of this modification also in *P. falciparum*. The non detection of yW in the MS study may be a technical issue of sensitivity, since we consistently detected low abundance of tRNA^Phe^ in *P. falciparum* using tRNAseq (Li et al. 2024).

During the asexual replicative cycle, we observed a downregulation of most detected modifications when the parasites transit from ring to trophozoite stage. Concurrently, we noted an increased level of intron-retaining precursor tRNA^Tyr^ during this stage. Pre-tRNAs can be used to assess the transcriptional activities of RNA Polymerase III which cannot be reliably determined by measuring the level of equilibrium mature tRNAs, which are often rapidly processed and have long half-live (Karkusiewicz et al. 2011; McLean and Jacobs-Lorena 2017; Shetty et al. 2020). The increase in pre-tRNA^Tyr^ is in line with the general increase of transcriptional activity in metabolically active trophozoite stage, and consistent with bulk maturation of tRNAs needed for the ramping up the translational activities. The detected modifications found in pre-tRNA^Tyr^ are unanimously lower when compared to mature tRNA^Tyr^, particularly for m^1^G37, which was entirely absent in pre-tRNA^Tyr^. This is consistent with finding in other organisms that tRNA guanine(37)-N1-methyltransferase can only recognize spliced tRNAs as substrate (Ohira and Suzuki 2011; Hopper and Nostramo 2019). Collectively, our results suggest that the lower modification levels observed during the trophozoite stage are likely attributed to an increased synthesis of tRNAs that creates a backlog of intermediately processed tRNAs, than an *ad hoc* stage-specific regulation of the epitranscriptome.

Our conclusion that there is no stage-specific regulation of the tRNA epitranscriptome is in apparent contradiction with the previous MS study which described a significant increase of many types of modification (22 out of 28) correlating to the parasite’s maturation. However, it is important to note that our methods do not present the same type of data. The MS study presented the relative ratio of a modified nucleoside to the unmodified nucleoside, therefore, the increase in a particular type of modification could be due to the increased expression of specific tRNAs that contain the modification. Wherein, our study estimates the change in the stoichiometry of each modification at single-base resolution, albeit to a much-limited repertoire.

Sexual commitment and differentiation from the asexual stage is critical for the parasite to complete its transmission cycle. Thus, blocking this biological process has emerged as an appealing strategy for disease control. Most key factors identified for the process so far are regulators on the epigenetic and transcriptional levels such as several Ap2 transcription factors (Lopez-Rubio et al. 2009; Brancucci et al. 2014; Coleman et al. 2014; Kafsack et al. 2014; Sinha et al. 2014; Brancucci et al. 2017), but the regulation on the posttranscriptional and translational level is less clear. A previous study demonstrated that m^5^C38 protects tRNA^Asp(GTC)^ from degradation in nutrient-limiting condition, and mutant strain which lost PfDNMT2 mediated m^5^C38 exhibited marked increase in nutrient stress-triggered sexual commitment (Hammam et al. 2021a). Intriguingly, m^5^C methylation of mRNA by NSUN2 in has also been shown to promote mRNA stability and sexual conversion *P. falciparum* (Liu et al. 2022). However, the authors in that study did not assess the potential confounding effect of NSUN2 on the tRNA epitranscriptome, which are conventionally annotated as tRNA m^5^C methyltransferase. These studies implicate tRNA epitranscriptome in the regulation of sexual differentiation. In our study, we saw variations on several types of modification on some tRNAs during gametocytogenesis, ascertaining the potential involvement of the epitranscriptome in regulating the process. Further studies are needed to establish this on a mechanistic level.

Codon-biased translation regulated by tRNA modification, particularly U34 thiolation, has been shown to directly contribute to chemotherapy resistance in tumors. Similarly, artemisinin resistance in *P. falciparum* is also dependent on translatome reprogramming U34 thiolation (Small-Saunders et al. 2024b). Furthermore, transcriptomics analysis on large-scale population level also discovered increased expression of several tRNA-modifying genes in artemisinin resistance strains after exposure to artemisinin (Mok et al. 2021). Our results also implicate the involvement of tRNA modifications in the response to artemisinin exposure, as a significant proportion of positions showed an elevated stoichiometry in the aftermath of a short DHA assault. Functional studies will be needed to further exploit these as possible strategy counter the continued spread of artemisinin resistance.

## Materials and methods

### Parasite Culturing

Three *Plasmodium falciparum* lines were used in this study. NF54 and its transgenic line Pf-Peg4-tdTomato, which was used to induce gametocytes, as well as PF3D7_1248100 knockout strains in NF54 background.

All cultures were maintained in human O^+^ erythrocyte at 3% hematocrit in RPMI 1640 medium (Thermofisher Scientific), supplemented with 50ug/L gentamicin (Gibco), 2mM L-glutamine (Hyclone), AlbumaxII (Gibco) and hypoxanthine (50mg/L). To keep the microaerophilic environment, the culture flasks were maintained at 37°C on a 50-rpm shaker and constantly gassed with 90% N2, 5% O2, 5% CO2 gas mixture.

### Gametocytes induction

NF54-Peg4-tdTomato was used in this experiment to generate gametocytes (McLean et al. 2019). Prior to induction, synchronized cultures (∼1% parasitemia at 4% hematocrit) were maintained at 37°C in 96% N2, 1% O2, and 3% CO2. When the cultures reached 10-12% parasitemia, gametocytes induction was carried out following previously described protocol (Fivelman et al. 2007). For the isolation of gametocytes, N-acetylglucosamine (GlcNAc) was continuously added to the cultures from day 0 onward to inhibit any further growth of the asexual stage parasites.

### Sample collection

Parasites were synchronized by two consecutive Percoll/ sorbitol treatments before the start of the experiment.

The time series experiment with NF54 started with 30ml culture at 3% hematocrit and ∼5% parasitemia. For small RNA isolation, 10mL of culture was collected at each time point: Ring: 8-12 hpi, Trophozoite: 24-28 hpi, Schizont: 40-44 hpi.

For gametocyte samples, 10mL cultures were collected for small RNA isolation at each time point, including the trophozoite stage as a control, and on day-1, day1, 3, 5 and 7 post-gametocyte induction.

For the DHA experiments, three flasks of tightly synchronized young ring-stage parasite cultures (0-3hpi, with 3% hematocrit and ∼5% parasitemia were exposed to 10nM and 700nM DHA, respectively, and 0.1% DMSO as control for 6 hours, after which they were directly collected for RNA isolation.

### Small RNA isolation

Small RNAs(<200nt) were extracted directly from the pellets of parasitized erythrocytes utilizing the *mir*Vana miRNA isolation kit (Invitrogen), following the manufacture’s protocols. First, 600μl lysis buffer was added to 150μl infected RBCs and the mixture was vigorously vortexed to ensure complete cell lysis. Subsequently, 1/10 volume of miRNA Homogenate Additive was added to the cell lysates, followed by a 10-minute incubation on ice. An equal volume of Acid-Phenol (relative to the lysate volume before the addition of the miRNA Homogenate Additive) was then added, followed by vigorous vortex and centrifugation at 10,000 x g for 5 mins. The aqueous phase was transferred to a new tube. Next, 1/3 volume of 100% ethanol was added to the aqueous phase, and the mixture was passed through a glass-fiber filter to enrich small RNAs in the flow-through. An additional 2/3 volume of 100% ethanol was then added to the flow-through, which was passed through a second filter to immobilize the small RNAs. The column was sequentially washed once using miRNA wash solution1 and twice with solution2/3. Finally, the small RNAs were eluted with 70μl pre-heated(95°C) nuclease-free water.

### Recombinant synthesis of AlkB demethylases

pET30a-AlkB and pET30a-AlkB-D135S were gifts from Tao Pan (Addgene plasmid # 79050 and #79051). These plasmids contain wildtype *E.coli alkB* or D135S mutant sequences, both with a deletion of 11 residues from the N-terminal (Cozen et al. 2015). Plasmids were transformed into *E.coli* BL21(DE3) T1R pRARE2 and the cultures were grown in the presence of 50 μg/ml kanamycin and 34 μg/ml chloramphenicol at 37°C until OD600 reached 2. Protein expression was then induced with 0.5 mM IPTG at 18°C overnight. Cells were pelleted and resuspended in lysis buffer (100 mM HEPES, 500 mM NaCl, 10% glycerol, 10 mM imidazole, 0.5 mM TCEP, protease inhibitor cocktail (Roche), pH 8.0). The Lysate was sonicated by pulsed sonication (4s/4s on/off cycle for 4 min, 80% amplitude) and then centrifuged and filtered through 0.45 μm filters. The soluble proteins were purified first using HisTrap HP (GE Healthcare) and then by gel filtration on a HiLoad 16/60 Superdex 200 column (GE Healthcare) on an ÄKTA Xpress system. Thrombin was added (1 unit/mg protein) to the purified recombinant proteins and incubated overnight at 4°C. The cleavage tag was then moved by HisTrap HP. The proteins were then concentrated in storage buffer (20 mM HEPES, 300 mM NaCl, 10% glycerol, 2 mM TCEP, pH 7.5).

### tRNA demethylation reaction

Demethylation assay was performed to validation the identities of the detected mutation signatures. 1 μg small RNA was treated with 80 pmol WT and 160 pmol D135S AlkB in demethylase reaction buffer (300 mM KCl, 50 mM MES buffer (pH 5.0), 2 mM MgCl2, 50 μM of (NH4)2Fe(SO4)2·6H2O, 300 μM 2-ketoglutarate (2-KG), 2 mM L-ascorbic acid, 50 μg/ml BSA). The reaction was incubated at room temperature for 2h and then quenched by adding 5 mM EDTA. The treated RNA was purified using RNA clean-up and concentration micro-elute kit.

### Bisulfite conversion

Bisulfite conversion of RNA was used to detect m^5^C methylation on tRNA. 250ng of small RNA was used for the reaction using EZ RNA methylation kit according to the manufacturer’s instructions.

### tRNA library construction

This procedure was performed using a modified protocol from the manufacturer’s instruction for TGIRT library construction (InGex, Inc).

For template/primer annealing, a 20μl reaction volume containing the annealing buffer (100mM Tris-HCl, pH 7.5, 0.5mM EDTA), R2R DNA and its complementary R2 RNA was incubated at 82°C for 2min, followed by gradual cooling down to 25°C with 0.1°C s^-1^. Template switching was achieved using TGIRT-III RT (InGex, Inc). Briefly, a total of 100ng tRNA was combined with 4pmol primer mixture. Pre-incubate tRNA/primer at room temperature for 30min in low salt buffer (50mM Tris-HCl, pH7.5, 75mM KCl, 3mM MgCl2), 5mM DDT and 500nM TGIRT-III RT and then add dNTPs to a final concentration of 1mM to initiate the reaction. The template switching reaction was carried out at 42°C for 16h. To terminate reaction, 1μl 5M NaOH was added followed by incubation at 95°C for 3min, neutralizing with 1μl 5M HCl. The reaction products were then purified by MinElute Reaction Cleanup Kit (QIAGEN 28204). The purified cDNA was ligated to previously adenylated R1R DNA (100μM R1R DNA containing UMI sequences was adenylated with 1mM ATP in 10x reaction buffer with Mth RNA ligase at 65°C for 1h and inactivated at 85°C for 5min) at 65°C for 2h. After inactivation at 90°C for 3min, the ligated cDNAs were purified with MinElute Reaction Cleanup Kit (QIAGEN 28204). Amplification was performed using 2X Phusion High-Fidelity PCR Master Mix (Thermofisher) with initial denaturation, followed by 12 cycles of 98°C 5s, 60°C 10s, 72°C 10s. The PCR products were cleaned with 1.3X Agencourt AMPure XP beads (Beckman A63880) to get rid of primer dimer products. Quantification of libraries was performed by Collibri Library Quantification Kit (Invitrogen) and quality was assessed on the Bioanalyzer with a High Sensitivity DNA analysis Kit (Agilent 5067-4626).

*R2 RNA:*

5’-GAUCGGAAGAGCACACGUCUGAACUCCAGUCAC-3SpC3/

*R2R DNA:*

5’-GTGACTGGAGTTCAGACGTGTGCTCTTCCGATCT

*R1R DNA:*

5’-/5Phos/GATANNNNNNNGATCGGAAGAGCGTCGTGTAGGGAAAGAGTGT/3SpC3/

*R1 multiplex PCR primer:*

5’-AATGATACGGCGACCACCGAGATCTACACTCTTTCCCTACACGACGCTCTTCC GATCT

*Barcoded primer:*

5’-CAAGCAGAAGACGGCATACGAGAT*[BARCODE]*GTGACTGGAGTTCAGACGTGTGCTC TTCCGATCT

### tRNA library quantification and statistical analysis

#### Read preprocessing

All tRNA-seq libraries were sequenced as single-end reads for 120 reads on an Illumina Nextseq 550 platform. A standard quality control was performed via FastQC (v.074) after base calling and demultiplexing. Prior to mapping, unique molecular identifiers (UMI) were extracted using UMI extract tools (v1.1.2). Reads were processed using Clip tools (v1.0.3) with a size selection to filter reads<15bp to remove adapter sequence and low-quality sequence. In our library, we anticipate that the tRNA transcriptome of the parasite will produce most of the sequencing reads, with the remaining reads originating from either other small RNA species or human tRNA transcriptome. To accommodate this, filtered reads were then aligned to a curated tRNA reference which included all *Plasmodium* nuclear and apicoplast tRNAs along with all unique human tRNAs from GtRNAdb using Bowtie2 (v2.2.1) with sensitive local mode. The reference curation process involved re-defining the exact 5’ and 3’ positions of all tRNAs, adding 3’-CCA tail to all tRNAs, changing the reference from A to G for those tRNAs with inosine modification at position 34, and adding a 5’G base to His-tRNA. After alignment, de-duplication was performed using UMI deduplicate tools (v1.1.2). Unaligned reads were subsequently aligned to the *Plasmodium* 3D7 genome (v. PlasmoDB-46) in sensitive local mode to identify mappings to other small RNA species. Two additional steps were performed to evaluate the mapping outcome. First, unaligned reads from the genome mapping were re-aligned to the tRNA reference in the very sensitive local mode, which should yield very few reads. Second, unaligned reads with 3’-CCA tail were randomly assessed by blasting to the Plasmodium 3D7 genome to ensure that most reads with 3’-CCA were mapped in prior operations.

These analyses were conducted using the Galaxy EU platform.

Fractions of mutation/truncation signature at position n was calculated as 1-(coverage of reference sequence at position n/ total coverage at position n+1). The reads at each position are obtained by either IGV or Freebayes tools (v.1.3.6)

### Plasmid construction and parasite transfection

Plasmid construction and parasite transfection were performed according to established protocol (Ghorbal et al. 2014).

The pUF1-*Cas9* and pL6-*egfp* plasmids were generous gift from Dr. Jose-Juan Lopez-Rubio.

The pL6-*PF3D7_1248100* plasmid was generated by modifying pL6-*egfp* plasmid, which contains sgRNA and donor DNA template for homologous recombination repair.

To construct pL6-*PF3D7_1248100* plasmid, the *egfp* homology regions in the pL6 plasmid were digested using the restriction enzymes EcoRI/NcoI and AflII/SpeI, and then replaced with *PF3D7_1248100* homology regions. Two fragments of the *PF3D7_1248100* homology regions were amplified using primers P1/P2 and P3/P4, each containing a 15-bp homology for InFusion cloning (as detailed in the Supplementary table 1). The sgRNA cloning oligonucleotides P5/P6, P7/P8 and P9/P10 included 20-bp guide RNAs flanked by 15-bp homology regions suitable for InFusion cloning. To insert sgRNAs, pL6-*PF3D7_1248100* was digested with the restriction enzyme BtgzI, followed by InFusion cloning. Sanger sequencing confirmed the successful incorporation of the *PF3D7_1248100* homology regions and sgRNAs.

NF54 strain was transfected with previously described by electroporating ring-stage parasites with linearized pL6-*PF3D7_1248100* plasmid and circular pUF1-*Cas9* plasmid.

pL6-*PF3D7_1248100* plasmid was linearized with restriction enzyme ScaI and purified by ethanol precipitation. 10ug linearized DNA and 50ug circular pUF1-*Cas9* were co-transfected into parasites. Pyrimethamine selection (50ng/mL) was initiated 6 hours post-transfection. Keep parasite under pyrimethamine pressure by changing medium with pyrimethamine every day for the first week and every other day a week after. Pyrimethamine-resistant parasites were obtained within 3 weeks of transfection.

### Statistical analysis and figure generation

Two tailed student’s t-test, Manny-Whitney U test were used for statistical analyses wherever specified. A priori assumption of p < 0.05 was used for statistical significance. Levels of significance were denoted as *, **, ***,**** in respect to p < 0.05, p <0.01 and p < 0.001 and p < 0.0001

All statistical analysis and plots were generated with R (V.1.4.1103)

## Data availability

All tRNA sequencing datasets generated in this study have been deposited into the NCBI Gene Expression Omnibus database under accession number GSE226632. The *Plasmodium 3D7* genome (v. *PlasmoDB-46*) can be accessed from plasmodb.org. This study did not generate new unique materials.

## Acknowledgement

We express our gratitude to the members of the Larsson laboratory and the Ribacke laboratory for their valuable input and feedback. Special thanks to Pan Tao and Addgene for providing the demethylase plasmids, and to E.Strandback, H.A.Korsah, and T.Nyman at the Protein Science facility of Karolinska Institutet for their assistance in synthesizing the recombinant proteins. We also acknowledge the support of the Freiburg Galaxy Team, funded by the Collaborative Research Centre 992 Medical Epigenetics (DFG grant SFB 992/1 2012) and the German Federal Ministry of Education and Research BMBF grant 031 A538A de.NBI-RBC. This research was made possible through a grant from the Knut and Alice Wallenberg Foundation (2017.0055.) to M.W. and U.R., a grant from the Swedish Research Council (2021-03141) to S.CL.C. and M.W., and a grant from Tore Nilsons Stiftelse För Medicinsk Forskning (2017-00532) to S.CL.C. S.CL.C. was also supported by a postdoctoral fellowship grant from Svenska Sällskapet för Medicinsk Forskning, while Q.L. received support from the Chinese Scholarship Council and a doctoral grant (KID #2018-01037) from Karolinska Institutet.

---

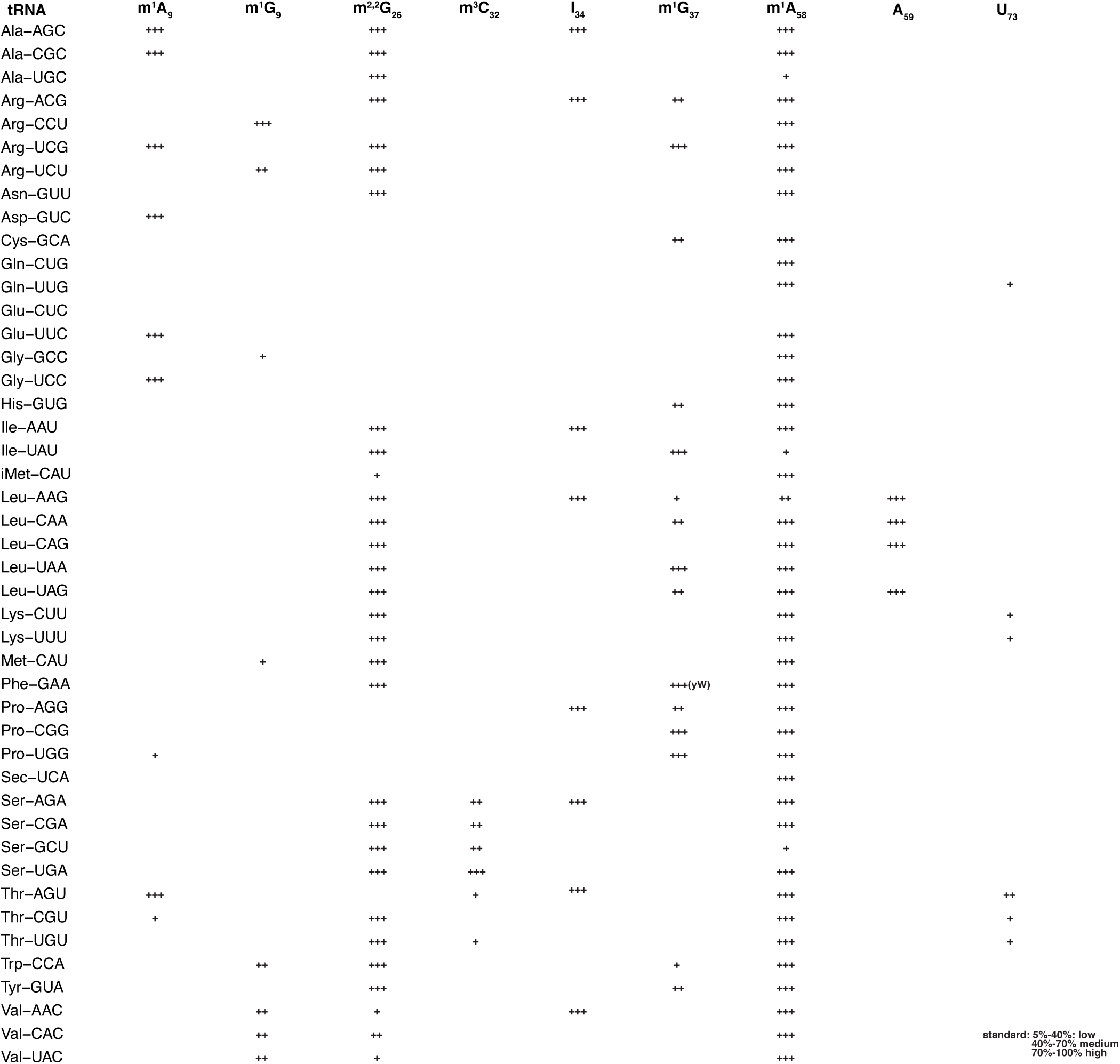

