## Supplementary figure 1-6 and supplementary table 1 for "Genome-wide profiling of tRNA modifications reveal the presence of divergent signatures in *Plasmodium falciparum*"

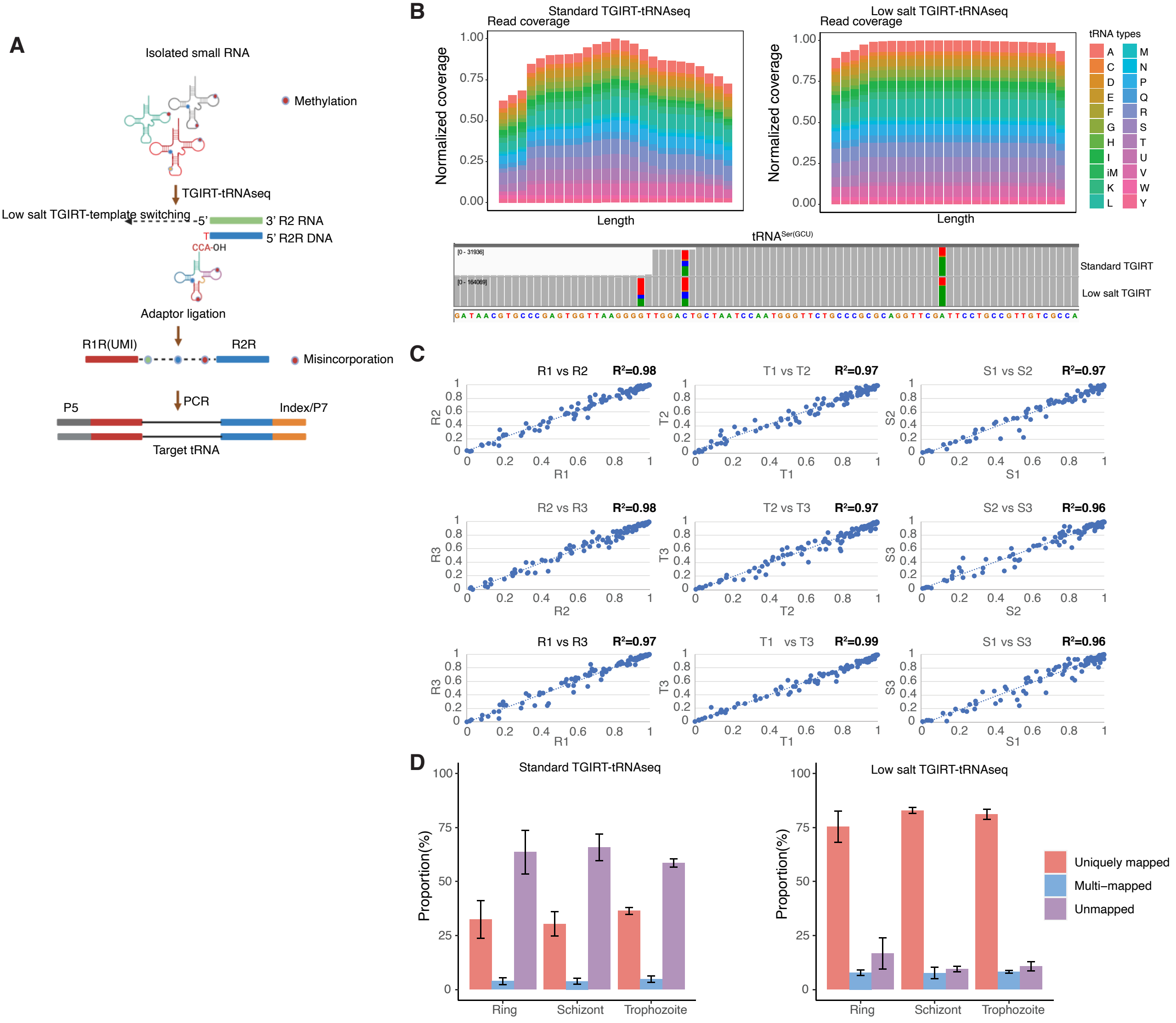
A
Isolated small RNA
Methylation
TGIRT-tRNAseq
Low salt TGIRT-template switching
5'
3' R2 RNA
5' R2R DNA
T
CCA-OH
Adaptor ligation
R1R(UMI)
R2R
Misincorporation
PCR
P5
Index/P7
Target tRNA
B
Standard TGIRT-tRNAseq
Read coverage
Normalized coverage
Length
Low salt TGIRT-tRNAseq
tRNA types
A C D E F G H I iM K L M N P Q R S T U V W Y
tRNASer(GCU)
Standard TGIRT
Low salt TGIRT
C
R1 vs R2 R²=0.98
T1 vs T2 R²=0.97
S1 vs S2 R²=0.97
R2 vs R3 R²=0.98
T2 vs T3 R²=0.97
S2 vs S3 R²=0.96
R1 vs R3 R²=0.97
T1 vs T3 R²=0.99
S1 vs S3 R²=0.96
D
Standard TGIRT-tRNAseq
Low salt TGIRT-tRNAseq
Proportion(%)
Ring
Schizont
Trophozoite
Uniquely mapped
Multi–mapped
Unmapped

| Stage | Category | Proportion(mean) | SD |
|---|---|---|---|
| Ring | P.falciparum nuclear-encoded tRNA | 0.867 | 0.012 |
| | P.falciparum apicoplast-encoded tRNA | 0.045 | 0.0099 |
| | Human tRNA | 0.087 | 0.021 |
| Trophozoite | P.falciparum nuclear-encoded tRNA | 0.954 | 0.005 |
| | P.falciparum apicoplast-encoded tRNA | 0.016 | 0.0061 |
| | Human tRNA | 0.030 | 0.010 |
| Schizont | P.falciparum nuclear-encoded tRNA | 0.943 | 0.015 |
| | P.falciparum apicoplast-encoded tRNA | 0.048 | 0.016 |
| | Human tRNA | 0.009 | 0.002 |

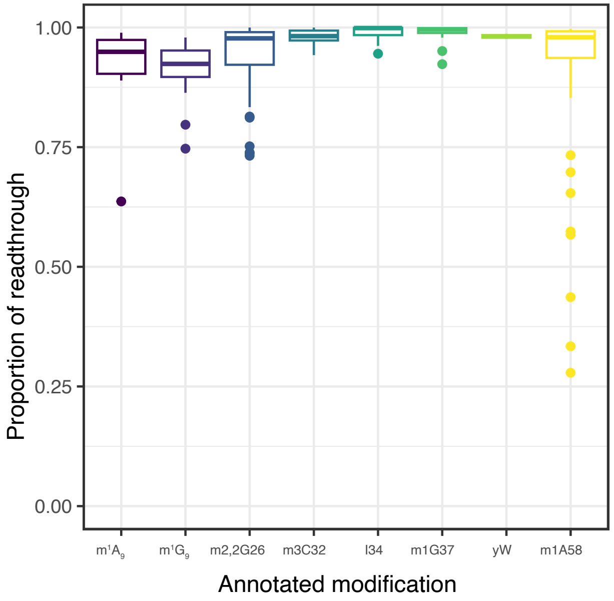

Proportion of readthrough
1.00
0.75
0.50
0.25
0.00
m1A9
m1G9
m2,2G26
m3C32
I34
m1G37
yW
m1A58
Annotated modification

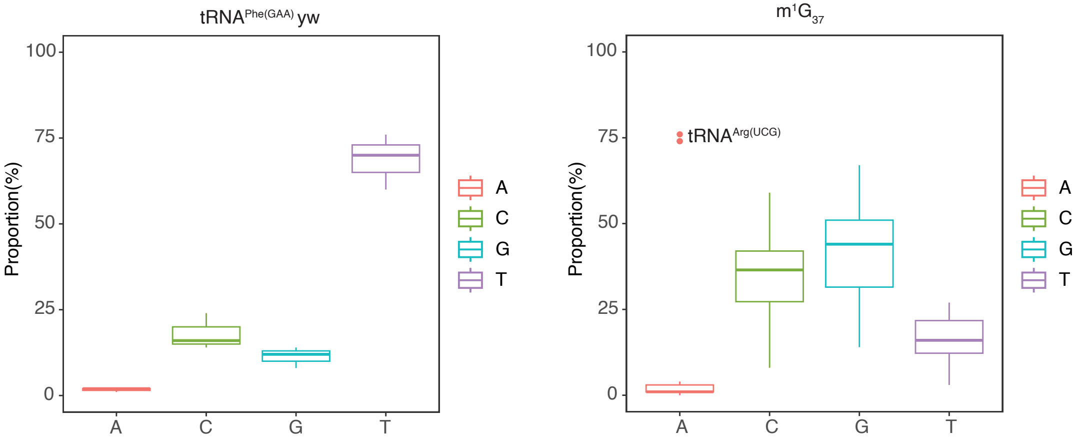

tRNAPhe(GAA) yw
m1G37
Proportion(%)
100
75
50
25
0
A
C
G
T
tRNAArg(UCG)

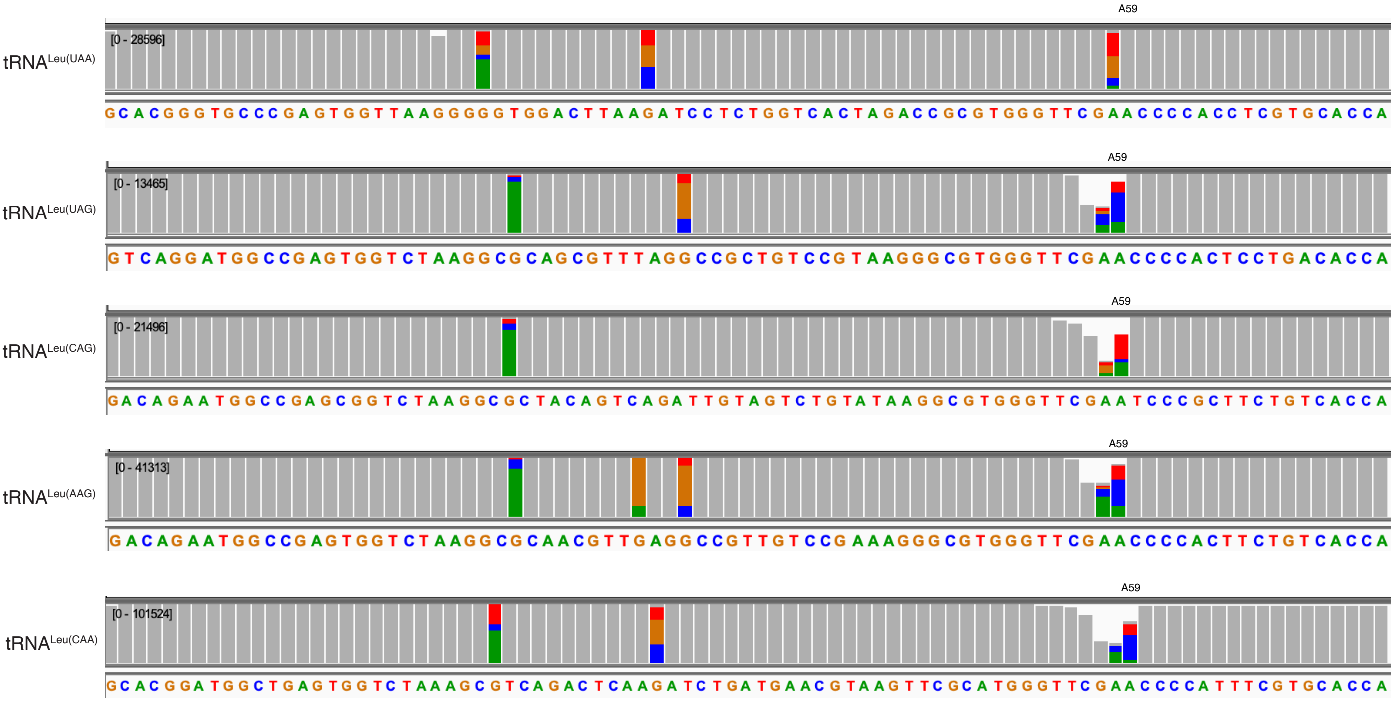
A59
tRNALeu(UAA)
[0 - 28596]
GCACGGGTGCCCGAGTGGTTAAGGGGGTGGACTTAAGATCCTCTGGTCACTAGACCGCGTGGGTTCGAACCCCACCTCGTGCACCA
A59
tRNALeu(UAG)
[0 - 13465]
GTCAGGATGGCCGAGTGGTCTAAGGCGCAGCGTTTAGGCCGCTGTCCGTAAGGGCGTGGGTTCGAACCCCACTCCTGACACCA
A59
tRNALeu(CAG)
[0 - 21496]
GACAGAATGGCCGAGCGGTCTAAGGCGCTACAGTCAGATTGTAGTCTGTATAAGGCGTGGGTTCGAATCCCGCTTCTGTCACCA
A59
tRNALeu(AAG)
[0 - 41313]
GACAGAATGGCCGAGTGGTCTAAGGCGCAACGTTGAGGCCGTTGTCCGAAAGGGCGTGGGTTCGAACCCCACTTCTGTCACCA
A59
tRNALeu(CAA)
[0 - 101524]
GCACGGATGGCTGAGTGGTCTAAAGCGTCAGACTCAAGATCTGATGAACGTAAGTTCGCATGGGTTCGAACCCCATTTCGTGCACCA

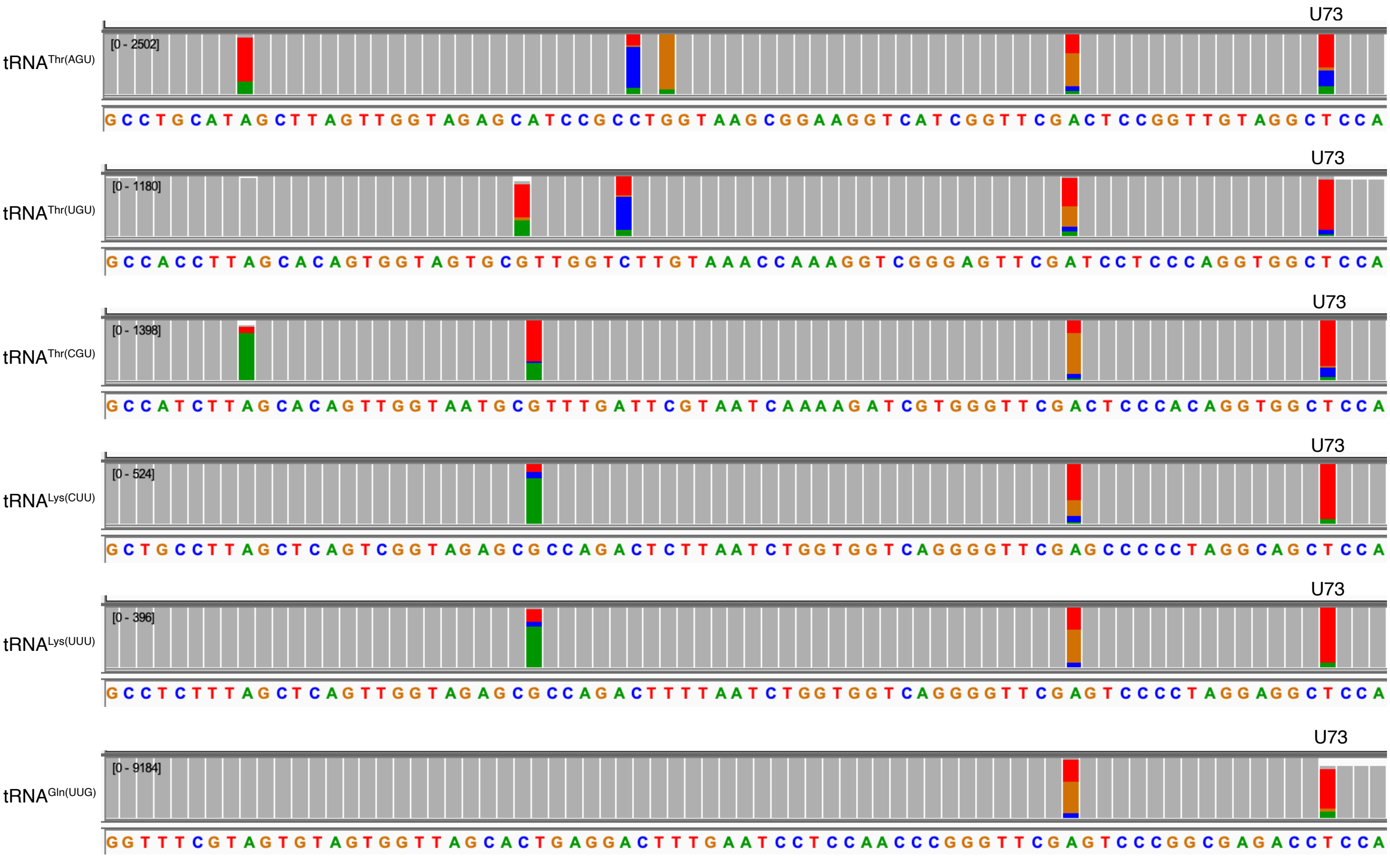

tRNAThr(AGU)
[0 - 2502]
U73
G C C T G C A T A G C T T A G T T G G T A G A G C A T C C G C C T G G T A A G C G G A A G G T C A T C G G T T C G A C T C C G G T T G T A G G C T C C A
tRNAThr(UGU)
[0 - 1180]
U73
G C C A C C T T A G C A C A G T G G T A G T G C G T T G G T C T T G T A A A C C A A A G G T C G G G A G T T C G A T C C T C C C A G G T G G C T C C A
tRNAThr(CGU)
[0 - 1398]
U73
G C C A T C T T A G C A C A G T T G G T A A T G C G T T T G A T T C G T A A T C A A A A G A T C G T G G G T T C G A C T C C C A C A G G T G G C T C C A
tRNALys(CUU)
[0 - 524]
U73
G C T G C C T T A G C T C A G T C G G T A G A G C G C C A G A C T C T T A A T C T G G T G G T C A G G G G T T C G A G C C C C C T A G G C A G C T C C A
tRNALys(UUU)
[0 - 396]
U73
G C C T C T T T A G C T C A G T T G G T A G A G C G C C A G A C T T T T A A T C T G G T G G T C A G G G G T T C G A G T C C C C T A G G A G G C T C C A
tRNAGln(UUG)
[0 - 9184]
U73
G G T T T C G T A G T G T A G T G G T T A G C A C T G A G G A C T T T G A A T C C T C C A A C C C G G G T T C G A G T C C C G G C G A G A C C T C C A

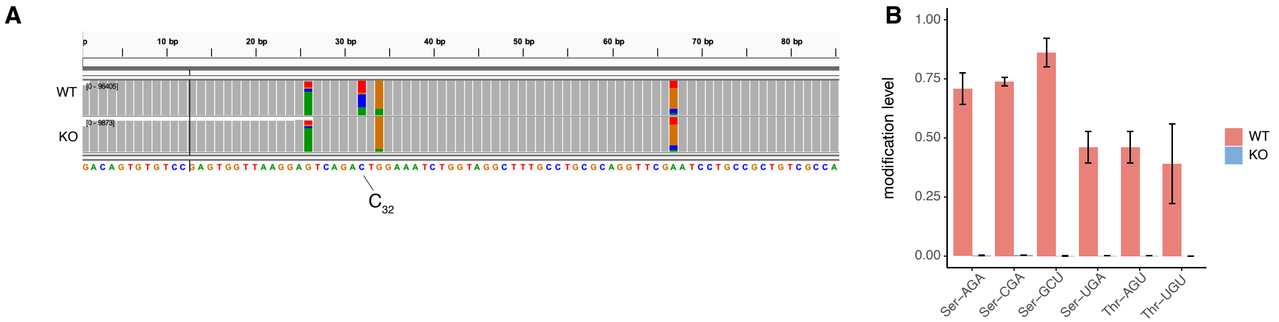
A
WT
KO
[0 - 96405]
[0 - 9873]
10 bp
20 bp
30 bp
40 bp
50 bp
60 bp
70 bp
80 bp
GACAGTGTGTCCGAGTGGTTAAGGAGTCAGACTGGAAATCTGGTAGGCTTTGCCTGCGCAGGTTCGAATCCTGCCGCTGTCGCCA
C32
B
modification level
1.00
0.75
0.50
0.25
0.00
Ser–AGA
Ser–CGA
Ser–GCU
Ser–UGA
Thr–AGU
Thr–UGU
WT
KO

### Supplementary figure legends

**Supplementary figure 1**
**Low salt TGIRT-tRNAseq significantly improves read coverage and the uniquely mapped ratio**
(A) Schematic showing tRNAseq library generation using TGIRT. A DNA/RNA duplex with a T overhang complementary to the terminal adenosine of mature tRNAs was used for the template-switching reaction. Modifications that interfere with the Watson-Crick pairing will cause misincoporation during reverse transcription. (B) Upper: Representative scaled 5'-to-3' sequence coverage plots spanning all nuclear tRNAs. The full-length tRNA was divided into 25 bins with each bin representing 4% of the tRNA length, as shown on the x-axis. The Y-axis value for each tRNA was normalized to the bin with the maximum coverage for each respective tRNA. Left panel: library generated using the standard protocol. Right panel: library generated using low-salt protocol. Lower: Example tracks for comparing the per-base read coverage along the full-length of Ser-GCU between libraries generated using the standard protocol and the low-salt protocol using IGVviewer. (C) Pairwise comparisons of the stiochiometry of all detected mutation signatures between biological replicates demonstrated high reproducibility. (D) Bowtie2 alignment-statistics for libraries generated using the standard (N=3) and the low-salt protocol (N=3). Error bars represent standard deviation.

**Supplementary table 1**
**Mapping statistics of tRNA libraries using low salt protocol**
Table showing the proportion of all reads mapped to tRNA genes base on their sequence origin. Mean and standard deviation (SD) are calculated froom N=3 biological replicates.

**Supplementary figure 2**
**Near complete readthrough of majority modifications enables the identification and quantification of modifications**
Boxplot showing the proportion of reverse transcription-readthrough per annotated modification. The mean readthrough proportion was calcluated from all detected mutation signatures of the same type modification on all nuclear-encoded tRNAs and from three biological replicates. The annotations were inferred *a priori.*

**Supplementary figure 3**
**The mutation signature at G37 of Phe-GAA**
Boxplots showing the base-called nucleotides identity due to RT misincorporation on two inferred modification sites; (left) yw (only on G37 of Phe-GAA) and (right) $m^1$G37.

**Supplementary figure 4**
**Leu tRNAs harbor mutation signature at A59**
Representative IGVviewer tracks showing the the per-base read coverage along the full-length of all $tRNA^{Leu}$. Mutation signatures were found on the A59 position of four out of the five $tRNA^{Leu}$. The threshold frequency for showing variant is 5% of the total called base.

**Supplementary figure 5**
**All $tRNA^{Lys}$, $tRNA^{Thr}$ and Gln-UUG harbor mutation signature at A59**

Representative IGVviewer tracks showing the the per-base read coverage along the full-length of all tRNAs harboring a mutation signature on the U73 position. The threshold frequency for showing variant is 5% of the total called base.

**Supplementary figure 6**

**PF3D7_1248100 is responsible for depositing $m^3C_{32}$ on $tRNA^{Thr}$ and $tRNA^{Ser}$**

(A) Representative IGVviewer tracks showing the the per-base read coverage along the full-length of Ser-AGA between wildtype and PF3D7_1248100 knockout parasites. Knocking out PF3D7_1248100 completely abolishes deposition of $m^3C32$ modification. The threshold frequency for showing variant is 5% of the total called base. (B) Bar graph showings the modification levels on all known $m^3C32$ sites between wildtype and PF3D7_1248100 knockout parasites
